# Narrative Context Shifts Gaze from Visual to Semantic Salience

**DOI:** 10.1101/2025.10.24.684352

**Authors:** Eva Berlot, Lea-Maria Schmitt, Christoph Huber-Huber, Marius V. Peelen, Floris P. de Lange

## Abstract

Humans make over a hundred thousand eye movements daily to gather visual information. But what determines where we look? Current computational models typically link gaze behaviour to visual features of isolated images, but we know that eye movements are also strongly shaped by cognitive goals: Observers gather information that helps them to understand, rather than just represent, the world. Within this framework, observers should focus more on information that updates one’s understanding of the environment, and less on what is purely visually salient. Here we tested this hypothesis using a free-viewing paradigm of narratives where we experimentally manipulated the meaningfulness of temporal context by either presenting pictures in a coherent, i.e. correct, order, or in a temporally shuffled order. We developed a novel approach to quantify which visual information is semantically salient (i.e., important for understanding): we separately obtained language narratives for images in stories, and computed the contextual surprisal of visual objects using a large language model. The ability of this semantic saliency model in explaining gaze behaviour was compared to a state-of-the art model of visual saliency (DeepGaze-II). We found that individuals looked relatively more often and more quickly at semantically salient objects when images were presented in coherent compared to shuffled order. In contrast, visual salience did not better account for gaze behaviour in coherent than shuffled order. These findings highlight how internal contextual models guide visual sampling and demonstrate that language models could offer a powerful tool for capturing gaze behavior in richer, meaningful settings.

## Introduction

We use visual information to build internal models of the world. However, at any given moment we only sample a small portion of our surroundings with high fidelity, as detailed visual input is limited to the fovea. As a result, we must move our eyes around to gather relevant details from the environment.^1,2^ But what determines where we look?

Eye movements are strongly influenced by physical properties of the visual input. We tend to fixate on informative or visually striking features, and avoid uninformative or homogeneous areas like blank walls. State-of-the-art computational models of visual sampling^3–5^ have formalized this notion by computing saliency maps of individual images. In these models, salience can be quantified from low-level visual features in images (e.g. luminance contrast, edge orientation^3,6^), features emergent in visual convolutional neural networks processing images,^4^ or local definitions of visual ‘meaningfulness’ of image patches.^5^ What all of these approaches have in common is that the salience is defined locally and quantified on the level of individual images. In the current paper, we use the term visual salience to refer to the output of these models (rather than to the specific visual features themselves, such as contrast values).

It is well known, however, that eye movements are also strongly shaped by top-down influences such as behavioral goals and instructions.^7–9^ Visual sampling varies depending on specific instructions participants are given; for instance, eye movements differ if participants are asked to estimate the age or wealth of individuals depicted in a painting.^7^ Yet in everyday life, there is usually no experimenter instructing us on our ‘task’; we are often engaged in open-ended and unconstrained activities without a specific goal – such as when watching a movie or visiting a gallery. Understanding what guides our eye movements in such naturalistic settings remains a major challenge. There are two main reasons for this gap: (1) the difficulty in defining and quantifying goals during free viewing, and (2) the dominance of artificial paradigms using isolated images that limit the possible influence of broader, especially temporal, context on viewing behaviour.

One compelling proposal regarding goals in natural settings is that humans are inherently driven to understand their environment by constructing internal models of the world.^10^ A growing body of evidence suggests that the brain assigns intrinsic reward to learning new information,^11–14^ and within this framework, eye movements help reduce model uncertainty by sampling information that updates our mental models.^14–16^ Thus, when an image is presented in isolation, eye movements may be drawn to its visually salient elements as the observer attempts to “understand” the image by focusing on areas rich in visual information. However, when embedded within a meaningful temporal context, eye movements may shift away from what is visually salient, and instead prioritize elements that are most informative for understanding the broader narrative.^17^ For example, a plain closed door in a film scene may attract attention not because it is visually striking, but because it holds narrative significance within the unfolding story. On this view, eye movements may be primarily in the service of understanding the ’bigger picture’, rather than simply parsing the visual input.

A useful experimental approach to study the influence of the broader, temporal context on viewing behaviour has been used substantially in the field of narrative perception,^17–20^ where viewers are presented with images that either follow one after the other so that they build up a coherent narrative, or are shuffled in temporal order thus preventing narrative understanding. By experimentally manipulating the coherency of the broader context, this approach strikes a balance between experimental control and ecological validity (manipulating from a more ‘naturalistic’ coherent setting to a more ‘artificial’ shuffled setting).^21–23^ Using such paradigms, it has been demonstrated that some basic aspects of gaze behaviour differ depending on the temporal order, or coherency, of image presentation.^19,20^ A theoretical framework called The Scene Perception & Event Comprehension Theory (SPECT^17^) can account for these differences. SPECT describes from a theoretical perspective how perceptual processes and model construction are coordinated during visual narrative processing. Specifically, it proposes that viewing narratives is comprised of an interplay of front-end processes (such as information extraction through eye fixations) and back-end processes (construction of the event model, working and long-term episodic memory) that iteratively work together to build an event model of a narrative. This integrative framework can account for several reported findings on differences in eye movements depending on the temporal meaningfulness of stimuli. However, what is currently still missing is a more computationally specified approach linking the back-end processes, like model creation, to front-end processes of individual eye movements.

Here we investigated how visual sampling differs depending on the ability to construct an internal model of the narrative. Building on the body of literature on narrative perception, we experimentally manipulated the temporal context of narrative by presenting images across time so they together formed a coherent narrative, or temporally shuffled the order, thus preventing narrative construction. To better understand where individuals look in images to build the model of the narrative, we developed a new method that quantifies narrative-based semantic salience in images. We developed our novel metric of semantic salience by leveraging AI-powered language models to capture the narrative context of the images that participants observed. This definition of semantic salience is very different from other types of salience computations,^5,24,25^ as it considers narrative context which unfolds over time, as opposed to focusing solely on single images. Additionally, we defined also visual salience in images using state-of-the-art models^4^ of visual sampling. We hypothesized that gaze behaviour shifts from visual to semantic salience, when moving from images presented in a shuffled temporal order to images presented in coherent order where a story can be formed and understood.

We related visual and semantic salience in images to two aspects of gaze behaviour: sampling frequency (how much?) and latency (when?) of different objects being fixated. This builds also on a decades long question, of whether we look more, or also earlier at things that semantically violate our expectations.^26,27^ To preface the results, we observed, in line with our hypothesis, that in coherent contexts, gaze behaviour shifted away from visually salient elements towards those with high semantic salience. This demonstrates that gaze behaviour is guided not just by what we see, but by what we understand – and that language models can provide powerful tools for modeling semantic influences on vision in naturalistic settings.

## Methods

### Participants

Forty-nine healthy adult volunteers were recruited for the experiment. All participants received informed consent before the start of the experiment. The study was approved by the local ethics committee (CMO 2014/288; CMO Arnhem-Nijmegen, The Netherlands) and was conducted in compliance with these guidelines. A total of 42 participants (6 males, 36 females, mean age = 24 years, range 18-40 years) was included in the final sample. Two were excluded due to poor calibration signal with contact lenses, 1 due to problems with vision later discovered and four due to poor data quality (>25% of eye signal missing). All of the included participants had normal vision and reported no history of neurological or psychiatric disorders.

### Eye-tracking

An Eyelink-1000 eye-tracker (SR Research) was used to record eye movements at the frequency of 1000 Hz. For each participant, the dominant eye was determined (using the peek-in-the-hole test), and tracked throughout the experiment. Participants sat 80 cm from the monitor, and were positioned in a chin rest. A nine-point calibration was performed at the beginning of the experiment. At the beginning of each subsequent block, a fixation dot appeared to which participants were required to fixate. If the deviation exceeded 2 degrees of visual angle (dva), calibration was repeated before proceeding to story viewing.

### Experimental procedure

Participants viewed four stories from the Frog Stories collection, with each story composed of a total of 24 images.^28–31^ These stories have been widely used for psychological experiments, in relation to language acquisition in development (Frog Stories corpora^32^), and to investigate visual sampling,^33^ and also including eye-tracking.^20^ Here, this stimulus set was chosen as the images clearly construct a narrative with purely visual material, without using any text or language. For each participant, two of the stories were presented in the correct, i.e. coherent, order, and for the two other stories, the order was shuffled across all images of the story (**Figure 1**). This experimental manipulation of temporal order allowed us to examine how visual sampling behaviour depends on the narrative context – whether or not the story can be meaningfully constructed. The temporal order of images was shuffled in the same way across participants. Shuffling was done randomly with the restriction that no two adjacent images were allowed to remain adjacent in the shuffled order. The assignment of stories to temporal order (coherent or shuffled) was counterbalanced across participants. Additionally, the order of presentation of the four stories was also counterbalanced across participants. Participants received minimal instructions; simply to freely view the images. Each image was presented for 5 seconds and there was a 2-second inter-image-interval where a gray background was presented. Participants received a break after they finished with viewing of each story. After all four stories were presented, they were each presented twice more (for a total of 3 viewings per story). At the end of the viewing, participants were prompted to write down on the screen what they remembered for each of the four stories. They also completed the ‘Need for cognition’ questionnaire afterwards.^34^ In total, the experiment was completed within 90 min. For this manuscript, only data from the first viewing is analyzed. The re-viewing of the stories, story reconstruction or questionnaire data were not included.

**Figure 1.**
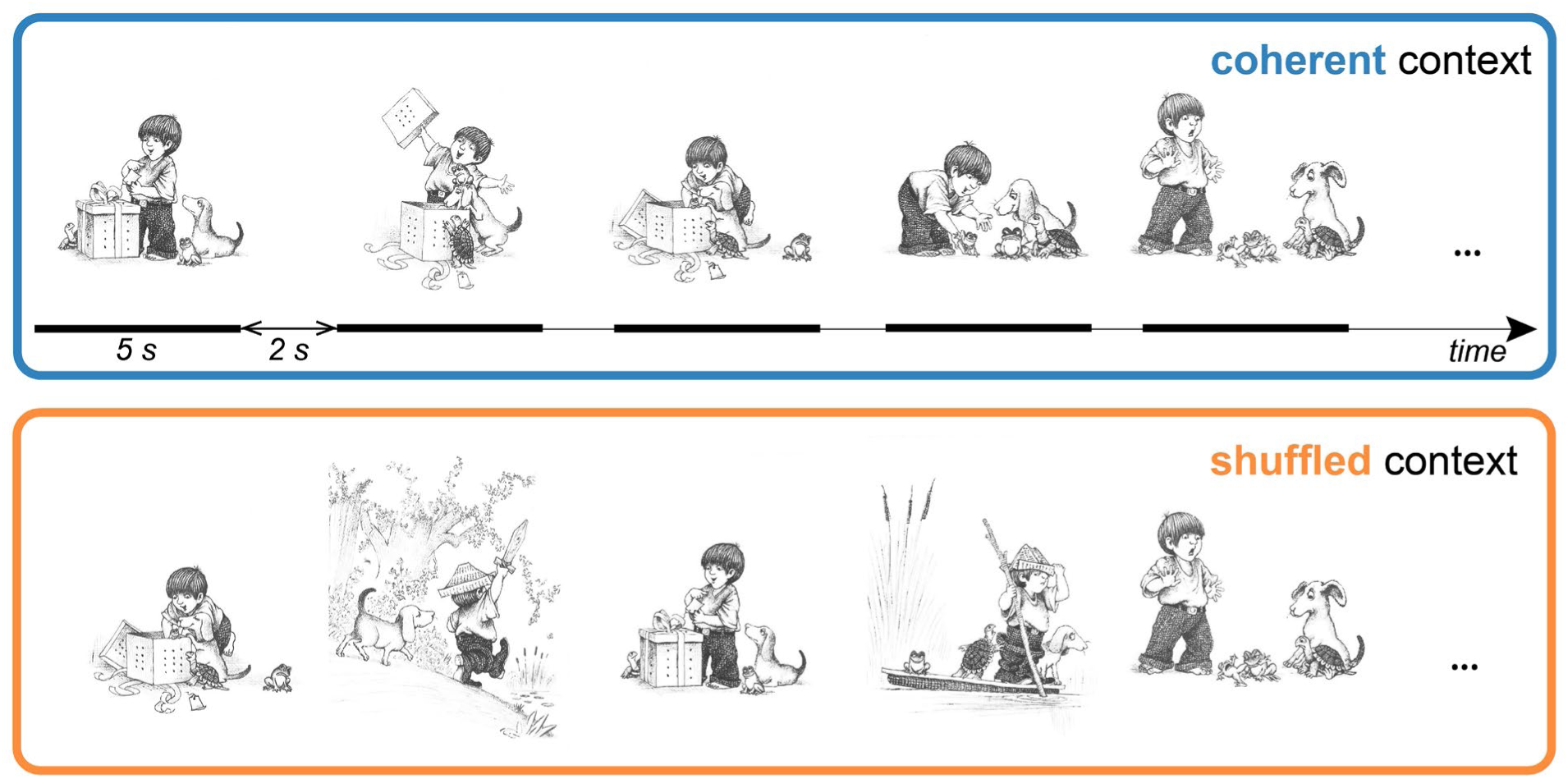
Depiction of images from the Frog Stories for one of the stories,^28^ as presented in the coherent (blue) or shuffled (orange) context. Each of the stories consisted of 24 images, here only 6 are depicted for illustration purposes. See Supplementary Figure 1 for depiction of all images in all four stories. Shuffling of images was performed within each story. Each image was presented individually for 5 s, with a 2-s blank interval between images.

### Defining objects

Objects in images were defined by drawing masks around them manually. Whenever objects were overlaid on top of each other, the resulting fixation was considered to be directed at the ‘top’ object (rather than the ones below it). On average, each image contained 7.2 objects (standard deviation of 2.3). The objects were, on average, 3.5 dva wide and 3.4 dva high. From the defined objects, 49% were animate (i.e. animals or humans), and 51% inanimate (scene elements such as trees, rivers, lake, or objects like gift box, boat etc.).

### Visual salience modelling

Visual salience was modelled by feeding the images to DeepGaze-II^4^, an eye-fixation prediction model that is based on a deep-neural network (VGG-19^35^) which is trained on object recognition, and additionally fine-tuned on human fixation data to predict image locations that attract most fixations (**Figure 2A-B**). It is the state-of-the-art model in performing human fixations, performing close to ceiling levels when applied to viewing behaviour of isolated images.^4^

**Figure 2.**
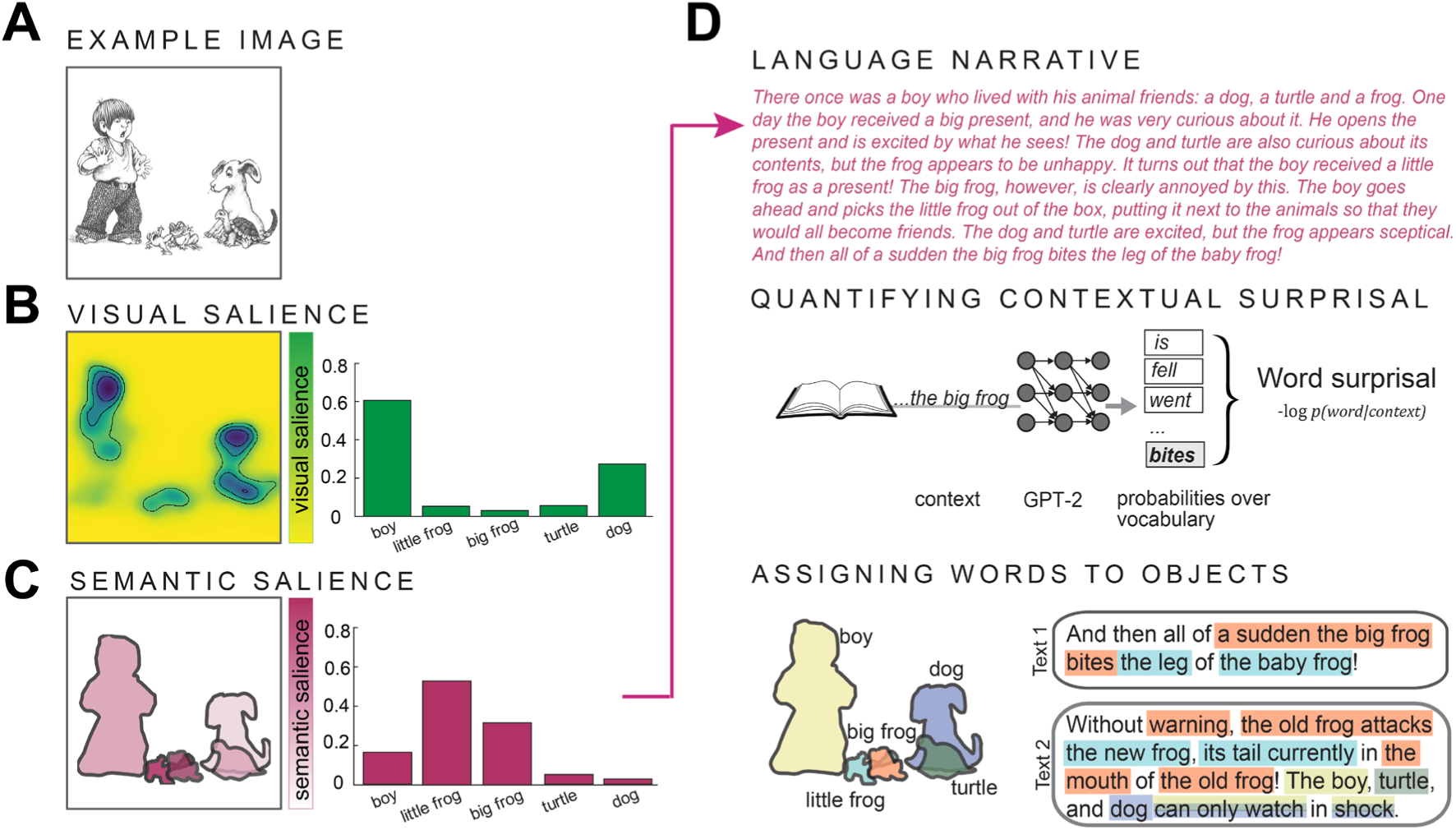
Modelling visual and semantic salience in images. **A)** Example image. **B)** Object-wise **visual salience** for the same image, extracted from DeepGaze-II.^4^ **C)** Object-wise **semantic salience** for the same image. **D)** Steps for establishing semantic salience. Narrative annotations were provided by an independent set of individuals (N=4). Using GPT-2,^37^ contextual surprisal was quantified for each word in the stories. Phrases of words (defined using Stanza^39^) were linked to objects and averaged across the four descriptions.

We also used a different salience model, Graph-Based Visual Salience,^6^ which is purely based on pre-defined image-computable features (i.e. not trained on human fixations), and thus does not exhibit any embedded cognitive biases that may be presented in DeepGaze-II by it having been trained on human fixation data. Therefore, it may be seen as a ‘purer’ estimate of image-computable visual salience. For both models, the object-wise values were extracted from each image. That was done as our new model of semantic salience (described below) assigns only object-wise salience level as opposed to pixel level. Therefore, this object-wise averaging allowed for a fairer comparison between the visual and semantic salience models. Values across objects were then normalized to sum to 1 in each image. Though our choice to focus on the object-wise level instead of considering the features within an object was primarily pragmatic so as to better compare the two models, note that there have been reports on some (central or preferred view) object locations being specifically important in determining salience.^36^

As both DeepGaze-II and GBVS exhibit central bias, i.e. assign higher probability of fixations ending in the centre of an image, we evaluated also a pure ‘central bias’ model to examine if any of the performance of the above-mentioned visual salience models may be purely attributable to central bias. To do so, we extracted the GBVS central bias to a uniform image that equated in size to our images. Then we applied this central bias image to objects in all our images to extract an index depending on central bias strength in an object’s location. As for the other models of saliency, we then normalized object-wise values to sum to 1 per image.

### Narrative descriptions

Four separate individuals, all native English speakers, provided narrative descriptions to images in English. They were instructed to write the descriptions accompanying each image, as they together made the coherent order (i.e. correct story order). As an example of image description, they were provided one image (not from the set they had to narrate) and a possible description. The descriptions per image were supposed to be not longer than a couple of sentences per image. For the rest, they were not given any specific instructions. Note that these individuals were not part of the main, eye-tracking, experiment. Additionally, our main experiment did not include these descriptions in any way. On average, the length of provided narratives was 652 words per story (644 for narrator 1, 543 for narrator 2, 700 for narrator 3 and 722 for narrator 4). Per image, the descriptions averaged to 27.2 words (standard deviation of 8.8 words).

### Narrative salience modelling

The obtained narratives were fed to GPT-2 (XL)^37^ (**Figure 2C-D**). GPT-2 is trained on a large amount of text to predict upcoming words, based on preceding context. Each story from each narrator was fed separately, whereby each token was predicted based on preceding context of up to 1024 tokens. The probability of the actual provided token was then read out and transformed into contextual surprisal values (-log(p)). Each words’ surprisal was normalized by its unigram surprisal (i.e. surprisal of the word without considering any context) value. This was performed to ensure that the surprisal value would not be driven primarily by word frequency of unexpected words, but truly reflect the *contextual* surprisal, i.e. in light of the preceding story context. The obtained word-wise surprisal metric reflects the degree to which a word deviates from expectations about the ongoing narrative, with higher values signaling more need for an update of the current mental model, thus more informational value.^38^ We then used Stanza^39^ to define how words were combined into phrases (**Figure 2D**). We used noun, verb, adjective and adverb phrases for subsequent mapping of text to objects in images. Two separate raters linked the phrases for each image’s description to objects in that image. A single phrase could be linked to more than one object (e.g. if it referred to an interaction between two characters). Afterwards, the discrepancies in assignment were discussed between the two raters to finalize the mapping of phrases to objects. Finally, the word-wise contextual surprisal values were averaged per object across words in a phrase. The values for all objects in an image were then normalized to add to 1. The resultant object-wise surprisal values were similar across the four narrators (*r*=0.69), and were then averaged across narrators to obtain one narrative salience value per object per image. This was performed also to ensure that various different linguistic structures and syntax across the four narratives would be averaged out, thus isolating the semantic, or meaning-based, aspect of salience for each object (**Figure 2D**). The obtained values of semantic salience were applied to modelling eye movements for both orders of story presentation (coherent, shuffled) to specifically capture how each element in the images emanates from the temporally evolving narrative. In this way, the model was also similar to the other visual salience models with each image having one quantification of visual salience (independent of condition, calculated by DeepGaze-II or GBVS), and one quantification of semantic salience (independent of condition, calculated on the basis of an independent set of narrators viewing coherently ordered image sequences).

### Data preprocessing

Fixations were defined as by default in the Eyelink online event parsing system, in which instantaneous thresholds of velocity (30°/s) and acceleration (8000°/s^2^) are used to determine the onset and offset of saccades; samples above threshold are determined to be in saccade, and samples below threshold are determined to be in fixation. The edf file from the EyeLink was imported using the EDFImport toolbox. Follow-up analyses on the eye-tracking data were performed using custom Matlab scripts.

#### Sampling frequency and latency

<u>The two main dependent metrics of eye movements were the sampling frequency and sampling latency. The two were defined for each object in each image. Specifically, sampling frequency was the number of fixations that landed per object during the 5 second viewing interval. Sampling latency was the timepoint of the first fixation that had landed on the object in an image. The two metrics were used in some subsequent analyses just for the most (visually or semantically) salient object, depending on the temporal order of presentation (coherent or shuffled order; **Figure 3**). In 25 out of 96 images, the most visually salient object was also the most semantically salient object. For the purposes of completeness, we left those images in the analyses though they did not allow differentiating between effects of semantic or visual salience. For other analyses,</u> every object’s sampling frequency and latency were considered and related to object-wise salience scores by computing Spearman correlations between the ranks of object salience and fixation frequency or latency (**Figure 5**).

**Figure 3.**
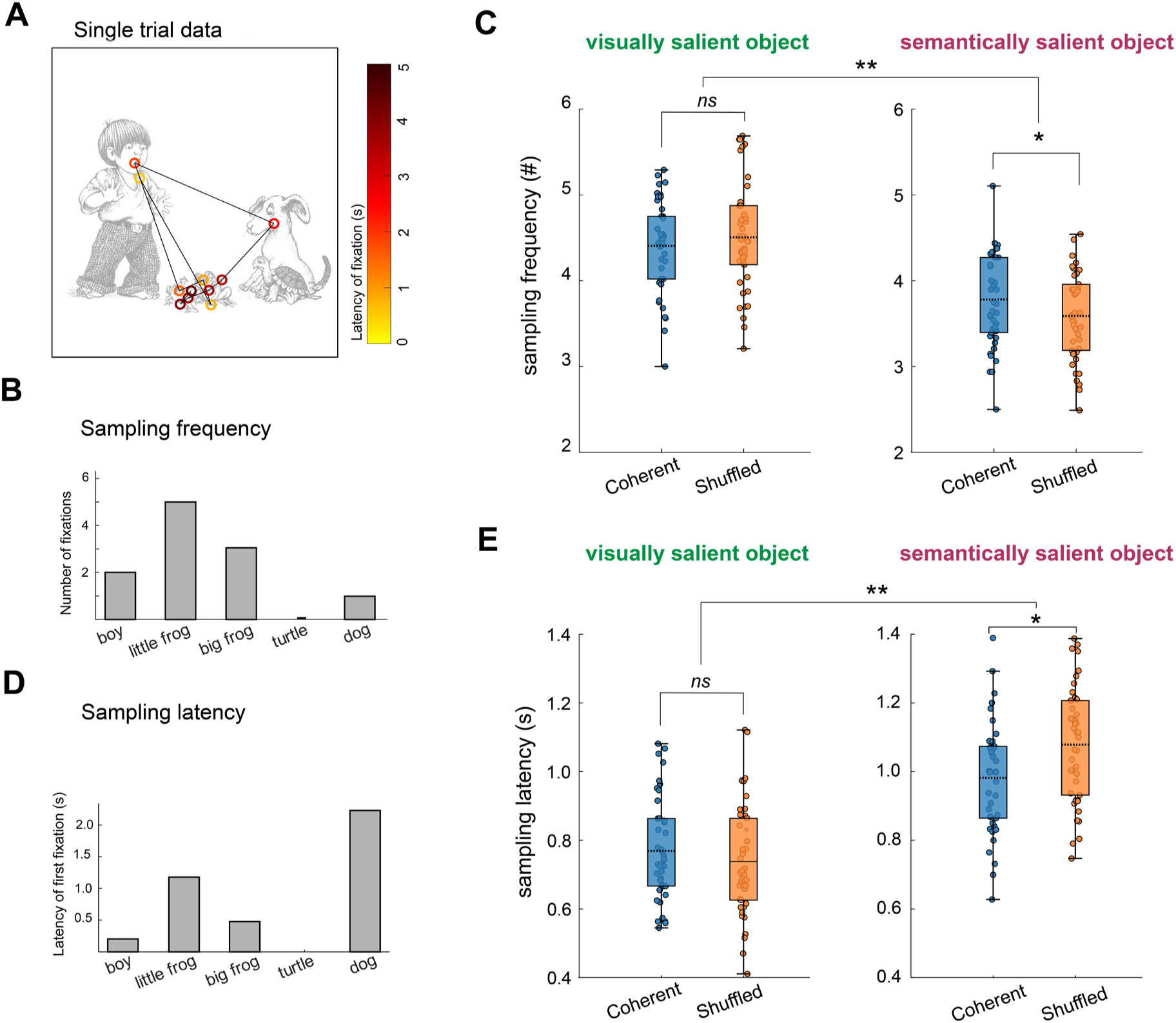
Sampling frequency and latency for visually and semantically most salient objects. **A)** Example single subject trial of one participant’s fixations. Colours indicate the latency of each fixation post-image presentation (yellow earlier than red). **B)** Sampling frequency extracted from A, i.e. fixation count per object. **C)** Sampling frequency of the most visually or semantically salient object across images, split by temporal order – coherent (blue) or shuffled (orange). * indicates significance level of *p*<.05, and ** *p*<.01, *ns* means not significant (*p*>.05). Individual dots represent individual participants – thus a total of 42 dots, one per participant in each condition subpanel. The central dotted line is the group mean value. Whiskers indicate values related to group deviation – the lower bound is quartile 1 – 1.5 x inter-quartile range and the upper bound quartile 3 + 1.5 x inter-quartile range. **D)** Sampling latency extracted from A, i.e. the latency of first fixation directed to each object in the image. **E)** Sampling latency of the most visually or semantically salient object across images, split by condition (coherent or shuffled order). Note that the relative sign of direction between coherent and shuffled conditions differs for results on sampling frequency and latency (i.e. coherent > shuffled for semantically salient object when examining sampling frequency, whereas coherent < shuffled for semantically salient object when examining sampling latency), as for frequency, more sampling leads to higher numbers, whereas for latency, earlier fixations lead to lower numbers.

### Probability of highest salience object fixation throughout the image presentation

The temporal evolution of sampling was assessed by testing the probability of each fixation directed to the most salient, visually or semantically, object for each ranked fixation during the 5-second image presentation, where 1 would be the first fixation after image onset, 2 the next etc. Specifically, we quantified for each fixation in a binary fashion whether the fixation was directed to the most (visually or semantically) salient object in the image. Across images for each participant, we then assessed the probability of each ranked fixation being directed to the most salient object, separately for coherent and shuffled orders. As number of fixations varied between images, we first considered (within subjects) fixations up to a reference value such that 75% of images had at least that many fixations. The same reference was applied also across subjects (75% of subjects having at least that many fixations), resulting in group analysis of 9 fixations per image (as plotted in **Figure 4**).

**Figure 4.**
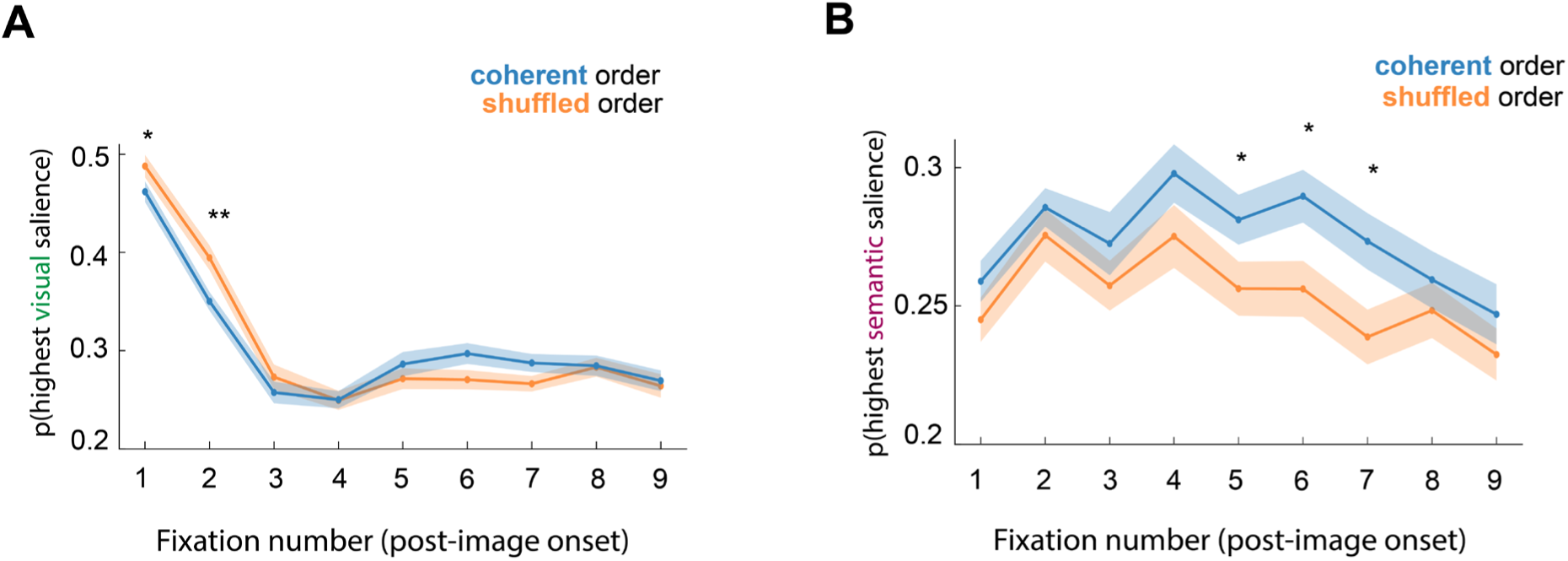
Sampling over the time course of image presentation. **A)** Probability of each fixation from image onset being directed to the most visually salient object. * indicates significance level of *p*<.05, and ** *p*<.01. **B)** Probability that each fixation from image onset is directed to the most semantically salient object.

#### Control analyses accounting for temporal order of image presentation

We accounted for two types of effects that could be driving some of the findings, both related to differences across subsequent images (i.e. images presented one after another). First we considered a possibility that any of the effects may be due to the most visually salient object staying in a more similar position across subsequent images for the coherent than the shuffled order. We computed the distance of the most visually salient object in each image compared to the one preceding image (depending on order) and included this as a covariate factor in a linear mixed effect model. Second, we considered the possibility that semantic salience may be related to the amount of pixel change across subsequent images may be related to semantic salience and therefore draw most visual attention – in other words, that participants may look for longer or earlier at the parts of an image with most pixel change across subsequent images. To account for that, we computed, for each most semantically salient object in an image, the amount of pixel change between subsequent images, and included this as a covariate in a linear mixed effect model.

### Statistical tests and two-sided reporting

Statistics for all of the reported tests are determined from a two-sided criterion (as opposed to one-sided or directed test). The manuscript and analyses were not preregistered.

## Results

Participants viewed images from picture stories, presented one at a time either in a coherent order or a shuffled temporal order, where the narrative structure of the images was disrupted (**Figure 1**). Each participant saw a total of four stories, two per condition (coherent or shuffled order), which allowed us to examine how each of the images was visually sampled depending on its narrative context—whether or not the story can be meaningfully constructed. We analyzed two key aspects of visual sampling, *sampling frequency* and *sampling latency*—i.e., how often and how quickly an object is fixated—to assess differences in overt sampling behaviour, depending on order coherence, and related those to visual and semantic salience of objects in images. Visual salience was defined using the state-of-the-art visual deep neural network DeepGaze-II^4^, which predicts human fixations based an image-computable features (**Figure 2A–B**; see Methods for details), whereas semantic salience was defined through a novel approach, using language modelling of surprisal through separately obtained narratives of picture stories (**Figure 2C-D**; see Methods for details).

### Stronger influence of semantic salience on sampling frequency in meaningful context

We hypothesized that gaze behavior shifts from being driven by visual salience to semantic salience, when participants view images in a coherent compared to shuffled temporal order. To test this hypothesis, we defined the most visually and semantically salient object in every image, and quantified how often these objects were fixated. An example is provided in **Figure 3A-B**. We expected that in coherent order, participants would preferentially sample the object most salient for narrative understanding, whereas in the shuffled order, they would preferentially sample the object that was most visually salient. In line with this hypothesis, we found a condition-specific effect of salience model (interaction between salience model and order condition: *F*_(1,41)_=12.570, *p*=9.96e^-4^). Specifically, semantically most salient objects were fixated more often when presented in the coherent, compared to shuffled temporal order (3.8 vs. 3.6 fixations; *t*_(41)_=2.54, *p*=.015). In contrast, there was no statistically significant difference in sampling frequency between conditions for the most visually salient object (4.4 vs. 4.5 fixations; *t*_(41)_=1.62, *p*=.113). In general, the most visually salient object was fixated more frequently than the most semantically salient object (4.5 vs 3.7 fixations; main effect of salience model: *F*_(1,41)_=190.98, *p*=5.0e^-17^; **Figure 3C**). Finally, there was no main effect of order condition on overall sampling frequency (*F*_(1,41)_<1). In sum, context influenced how much the different objects were sampled; when an image was embedded in a coherent temporal sequence, more eye movements were directed towards the most semantically salient object.

To rule out the possibility that adaptation influenced our results, we performed two control analyses. First, we considered the possibility that that participants were (marginally) less drawn to the most visually salient object in the meaningful condition because it remained in a similar position across neighboring images. To account for that, we included the distance between the centre of the visually most salient object in current compared to previously presented image as an image-specific covariate. The patterns of results remained unchanged (i.e. significant linear mixed model interaction salience type x condition: *F*_(1,7686)_=7.55, *p*=6.0e^-3^). Second, we considered the possibility that participants may be paying more attention to changing regions in the coherent compared to shuffled order because the objects in the image that change across subsequent images in a pixelwise fashion are also the ones that are considered more semantically salient. We included the amount of pixelwise change of the most semantically salient object as an image-specific covariate. The patterns of results also remained unchanged (i.e. significant linear mixed model interaction salience type x condition: *F*_(1,7351)_=8.35, *p*=3.9e^-3^).

On average, participants made more fixations in the shuffled than the coherent condition (15.3 vs. 14.9 fixations; *t*_(41)_=2.84, *p*=6.94e^-3^; see Supplementary Figure 1). To ensure that any of the observed effects were not driven by this absolute difference, we have repeated the analysis by considering the relative proportion, instead of absolute amount, of fixations. The relative proportion of fixations was defined as the ratio of number of fixations directed to the most visually or semantically salient object divided by the absolute number of fixations in that trial / for that image. The observed patterns of the results remained the same (see Supplementary Figure 2).

We further validated our findings using an alternative model of visual salience—Graph-Based Visual Salience (GBVS^6^) instead of DeepGaze-II (Supplementary Figure 3). GBVS offers a purer estimate of visual-only salience, as it relies solely on low-level, image-computable features, unlike DeepGaze-II which is additionally trained on human eye movement data which may introduce into the model other ‘non-visual’ factors and higher-order biases, all possibly making DeepGaze-II a ‘stronger’ model. Importantly, the context-specific effects of salience on fixation frequency were replicated with GBVS, thus supporting our central hypothesis that narrative context modulates the impact of salience on gaze (away from visual salience).

### Stronger influence of semantic salience on sampling latency in meaningful context

Having assessed *how much* individuals sample salient objects in images, we next turned to the question of *when* they first sample these objects (**Figure 3D**). Again, we observed a significant interaction between the salience type and temporal order in which images were presented (*F*_(1,41)_=5.95, *p*=.019), such that the semantically most salient objects were fixated earlier in coherent than shuffled order (0.98 vs. 1.04 s, difference of 96 msec; *t*_(41)_=2.98, *p*=6.46e^-3^). In contrast, the latency of fixation to the most visually salient object did not differ across conditions (0.77 s in coherent vs. 0.74 s in shuffled condition; *t*_(41)_=1.01, *p*=.32). The visually most salient object was overall fixated earlier than the semantically most salient object (0.75 s vs. 1.03 s post-image onset; ANOVA main effect of salience model, *F*_(1,41)_=123.51, *p*=6.05e^-14^; **Figure 3E**). Sampling latency itself did not differ significantly between conditions (coherent vs. shuffled; *F*_(1,41)_=3.13, *p*=.084). In sum, semantic salience influenced *when* attention was directed to objects—specifically, semantically salient objects were fixated earlier when participants could incorporate them in the unfolding story. Again, the context-specific modulation of salience was observed also when considering an alternative model of visual salience (Supplementary Figure 3C).

Additionally, just like for sampling frequency, also here we ruled out the possible effect of adaptation on participants being (marginally) earlier drawn to visually salient object in the meaningful condition by performing a covariate analysis, where the results remained unchanged (linear mixed model significant interaction salience type x condition: *F*_(1,6857)_=10.5, *p*=1.2e^-3^). Similarly, the interaction remained significant when considering the amount of pixel change as a possible covariate (linear mixed model significant interaction salience type x condition: *F*_(1,6564)_=8.35, *p*=3.0e^-3^).

### Visual and semantic salience affect distinct sampling windows

Next we wanted to delineate in more detail how the context of image presentation influences sampling behaviour as it evolves over time. We analyzed the proportion of times each sequential fixation (ranked from image onset) was directed to either the most visually or semantically salient object. This revealed a sharp drop in the likelihood that fixations were directed toward the most visually salient object after the first two fixations (**Figure 4A**; main effect of fixation number: *F*_(8,328)_=100.62, *p*=1.4e^-83^). This is in line with prior work showing that DeepGaze-II predicts worse fixations with higher ordinal number.^40^ Moreover, the first few early fixations were more likely to land on the visually salient object in the shuffled than in the coherent condition (ANOVA interaction fixation number x context: *F*_(8,328)_=2.55, *p*=.011). Furthermore, we repeated the analysis with GBVS which relies on more low-level visual features and observed a similar pattern of results (Supplementary Figure 5A).

To see whether this initial tendency of first few fixations being more likely directed to the most visually salient object is driven by both DeepGaze-II and GBVS’s central bias (i.e. initial fixations being more likely to be directed more centrally, and that more for shuffled than coherent context) we further repeated the analysis considering the pure ‘central tendency bias’ model, i.e. defining which object scores the highest in terms of potential central bias and assessing how likely individuals were to fixate on it with each fixation. We observed that although the first two fixations were overall more likely to be fixated at the highest center bias object, there was no statistically significant interaction with the temporal order (coherent or shuffled condition), as was observed above for the visual saliency models (ANOVA interaction fixation number x context: *F*_(8,328)_=0.53, *p*=.88). Therefore, the higher likelihood of sampling the most visual salient model in shuffled than coherent context is unlikely due to a central bias tendency.

Fixation behavior towards the most semantically salient object showed a different temporal profile. There was no sharp drop across fixation ranks, though there was a main effect of fixation number (*F*_(8,329)_=5.44, *p*=1.90e^-6^), with higher values for slightly later fixations. We observed also a main effect of context condition (*F*_(1,41)_=9.28, *p*=4.04e^-3^), with fixations in the coherent context being more likely towards the most semantically salient objects. This was specifically significant for fixations slightly later during image viewing (fixations 5-7), which were more likely directed to the most semantically salient object in the coherent compared to the shuffled context (**Figure 4B**). Together, these results reveal a striking temporal shift in salience-driven behavior: early fixations are strongly guided by visual salience, and more so when images do not make up a coherent narrative. In contrast, subsequent fixations are more often directed at the most semantically salient object, specifically when images are presented in the coherent temporal order.

### Temporal order influences sampling of all objects with different visual and semantic saliences

So far, we have focused on the most salient objects in images – those with highest visual or semantic salience – and how much visual attention they received. However, real-world environments and scenes usually contain many objects, and understanding gaze behaviour beyond the most salient ones is essential.

To assess whether our findings generalize to sampling of all objects, we tested for a monotonic relationship between object-wise sampling frequency and their visual or semantic salience. Specifically, we computed Spearman correlations between the ranks of object salience and fixation frequency, separately for coherent and shuffled orders (**Figure 5A**; i.e. where one data point corresponds to one participant value). Spearman’s correlation was preferred over Pearson’s as it does not assume normality or linearity, and works on our (salience-)ranked data. Mirroring our earlier results, we found that the relationship between object salience and fixation frequency depended on the condition (temporal order x salience type interaction: *F*_(1,41)_=17.12, *p*=1.7e^-4^). Semantic salience was more consistently associated with object sampling in the coherent order (*t*_(41)_=5.28, *p*=4.5e^-6^). The opposite was also true—visual salience and sampling frequency were more related in the shuffled order (*t*_(41)_=2.19, *p*=.035). This indicates that the association between context and type of salience shapes visual sampling across objects.

**Figure 5.**
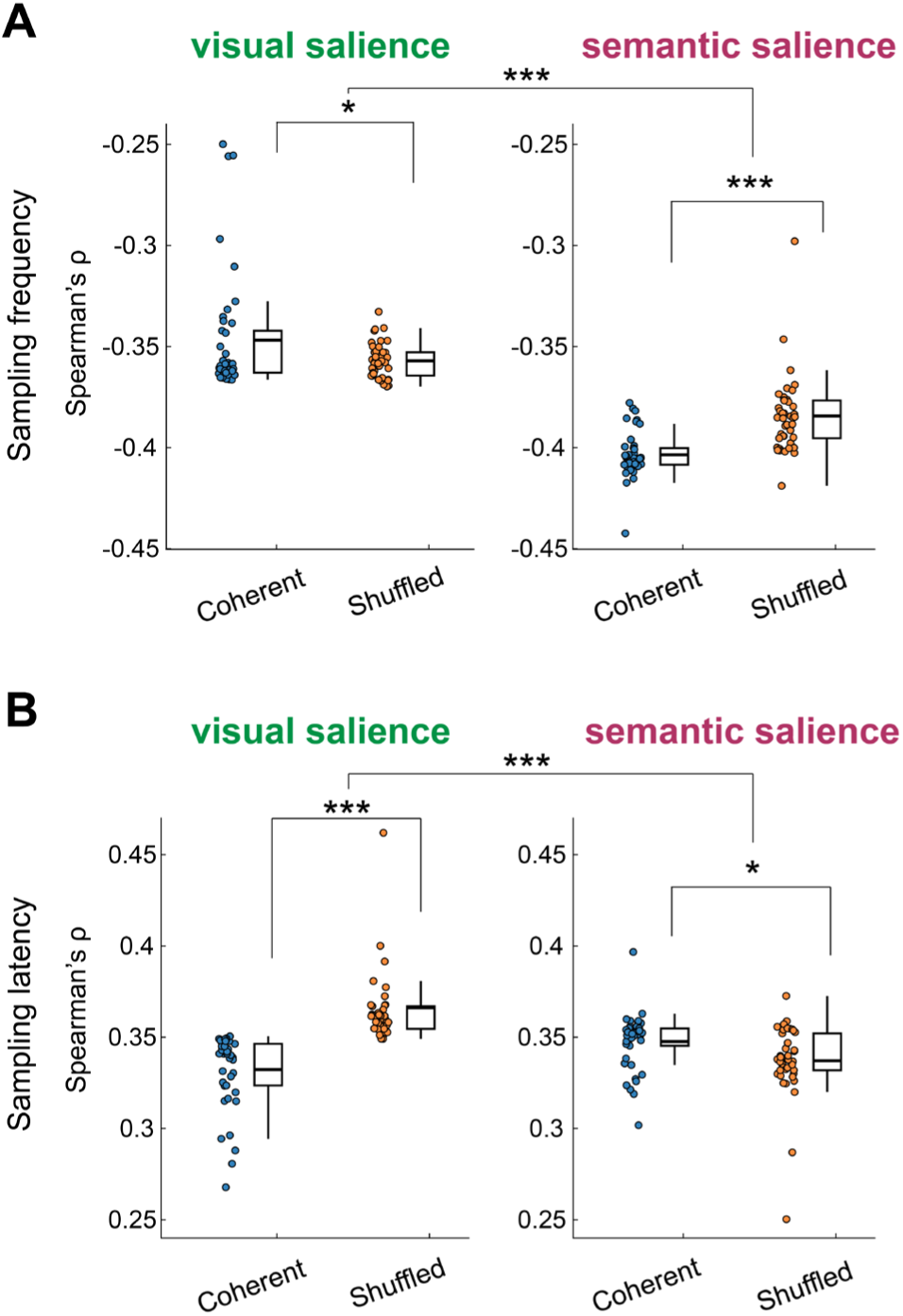
Salience influences sampling across all objects. **A)** Results showing the strength of a monotonic relationship between visual / semantic salience and sampling frequency, separated by condition (coherent vs. shuffled order). * indicates significance level of *p*<.05, ** *p*<.01 and *** *p*<.001. Note that the Spearman’s rho coefficients are negative because lower rank numbers correspond to higher salience (i.e., rank 1 is the most salient object), and therefore a negative correlation reflects a consistently positive mapping between higher salience and increased fixation frequency. **B)** Results showing the relationship between visual / semantic salience and sampling latency, split by condition. Note that now the sign of the Spearman’s rho coefficients is positive, reflecting that objects with higher (i.e., less salient) ranks are fixated later in time.

Similarly, we also examined whether there was a monotonic relationship between salience and sampling *latency* across all objects (**Figure 5B**). Consistent with the sampling frequency results, we observed a significant interaction between salience type and condition (*F*_(1,41)_=22.41, *p*=2.63e^-5^). Specifically, semantic salience was more reliably associated with fixation latency in the coherent order (*t*_(41)_=2.55, *p*=.014), whereas visual salience showed a more consistent monotonic relationship in the shuffled order (*t*_(41)_=4.85, *p*=1.83e^-5^). Similar results were observed, both for sampling frequency and latency, when considering GBVS as the salience model (Supplementary Figure 5A-B). Also for a model just considering the central tendency bias, a similar pattern of results was found (Supplementary Figure 5C-D), although the overall Spearman’s rho values were significantly lower for the central tendency model than for DeepGaze-II (fixation frequency: *t*_(41)_=126.42, *p*=9.1e^-55^; fixation latency: *t*_(41)_=192.22, *p*=3.2e^-62^).

## Discussion

We move our eyes to gather relevant visual details and build internal models of the world. But what determines where exactly we look, especially when there is no explicit task? We here built on ideas in visual neuroscience^10^ and narrative understanding^17,41^ fields, and in line of those hypothesized that an inherent goal of observers may be to form accurate internal models of the environment; and thus that eye movements are in the service of understanding, rather than just representing the visual environment. To investigate this, we used a free-viewing paradigm in which individuals viewed images presented in a sequence, where the temporal coherency of image order was manipulated–images were either presented so as to form a coherent narrative context over time or were shuffled in their temporal order. We found that this temporal, i.e. narrative, order systematically shaped visual sampling: in temporally coherent order, participants were more likely to fixate on objects that were semantically informative rather than those that were purely visually salient, both more frequently and earlier in time.

### Cognitive influences on gaze behaviour, as revealed by temporal order manipulation

Our findings demonstrate that eye movements are not merely reflexive responses to visually striking features, but are actively guided by viewer’s cognitive goals. Rather than passively reacting to the visual environment, individuals use their gaze to seek out information that helps reduce uncertainty related to their currently relevant internal models of the world.^14^ This highlights the fundamentally cognitive nature of eye movements and how they are sensitive to meaning, context, and the viewer’s evolving understanding. Our results align with the view that gaze serves as a tool for hypothesis testing,^10,13,16,42^ allowing observers to selectively sample information most relevant for constructing an accurate and coherent mental model. More specifically, our results are also in line with the SPECT framework, which describes how gaze behaviour is influenced by cognitive factors in the quest to build an event model of a narrative.^17^

Our manipulation of cognitive influences was by perturbing the temporal order in which images were presented so that in the coherent, i.e. correct, order viewers were able to construct a narrative, but this was prevented by shuffling the images. Previous studies have already shown that the temporal order has a strong influence on information sampling.^19,20,33^ Specifically, it has been shown that when images are presented in the way that participants cannot form a prior from preceding temporal context (somewhat analogous to our shuffled order), they exhibit more exploratory gaze behaviour,^19^ including making more fixations. Similarly, we here also observed that participants made more fixations when images were presented in the shuffled than coherent order. Thus this also likely reflects less of a need to visually explore images in coherent order due to a better formed prior from preceding temporal context.

Along the same lines, some of the previous studies that even used the same Frog picture stories as used here^20,33^ have also shown that when the image presentation order is perturbed, participants sample images for longer. In Hutson et al. (2018), it was specifically shown that this longer sampling was due to making 20% more fixations (as opposed to for instance fixating for longer) in the perturbed order. In all of these studies, the main finding can be attributed to viewers making an effort to find information that could help in understanding the narrative in the face of a lack of coherence. While in these two studies^20,33^ participants were left to determine sampling time themselves, in our study this was predetermined and fixed for them. Nonetheless, we also found an increased number of fixations in shuffled than coherent order. This aligns with the idea put forward by Hutson et al. (2018) that sampling more (rather than longer) may be a more effective strategy when faced with the lack of coherence or event model.

Interestingly, in the Hutson et al. (2018) study the authors found that this increase in the number of fixations was related to increased sampling of parts of the image that helped with story understanding. This differs from our finding that participants sampled the semantically most salient objects more in the intact than the shuffled condition. We believe that this difference in results may be attributed to differences in the design, specifically the coherency manipulation. Hutson et al. (2018) used a rather ‘local’ perturbation of image order – where the whole stories were broken up into local 3-image sequences that each consisted of a beginning state (e.g., the boy takes the little frog out of the box), a bridging event (e.g., the jealous big frog attacks the little frog), and an end state (e.g., boy picks up the little frog and scolds the big frog). Then, for the ‘incoherent’ presentation, the ‘bridging event’ was removed, and the authors compared the eye movements for the ‘end state’ image. Thus, this coherence manipulation was highly targeted and the observers could in principle still build a model of the story, and presumably did after fixating more on the parts of the image deemed relevant to bridge the event and complete the story. In contrast, in our study the coherence manipulation was more global (across all images), where images were throughout shuffled so that story understanding was (most likely) prevented. We believe that therefore participants were not able to determine what would have been semantically salient, and thus did not look at those objects as much. Both studies together, make an interesting new prediction – that for as long as the narrative model can be constructed (e.g. by having local perturbations of context) participants may look for *longer* at the semantically salient object, also as defined with our new metric. But once the coherency breaks down to the extent that such a model can no longer be formed, the sampling behaviour may be dominated more by visual saliency.

### The influence of semantic salience on eye movements takes time to build up

Our results of how visual and semantic salience operate over time bring new insights into a decades long question of how semantic information is processed in scenes. An early study suggested that observers fixate earlier, more often, and for longer on objects that seem semantically surprising in a scene (e.g. octopus in a farm scene) as opposed to objects that are likely to appear (e.g. tractor in a farm scene).^26^ They even found that the viewers were more likely to move their eyes to a semantically inconsistent object immediately, i.e. as the first fixation on the scene. Later studies have not replicated these early findings,^27^ especially the aspect that latency, and the earliest fixations may target the semantically inconsistent object. Since then, the dominant idea has been that when an image appears, attention is first guided by visual salience, with cognition playing a larger role as viewing unfolds^43,44^ (but see also ^25^). Our design adds a broader context of temporal order, which may provide additional help with early orientation and lead to earlier top-down influences. Nonetheless, we observed that the first few fixations are still much more likely to be driven to visually salient objects, even for the temporally coherent stories. Thus, the visual salience still dominates, before dropping off after the first few fixations. However, those first few fixations were less likely to be directed to the most visually salient object when images were presented in a temporally coherent order. Our control analyses showed that this was unlikely simply due to central fixation bias. Thus this may suggest that with temporal coherency, the visual salience is less dominant initially, but it nonetheless still takes a few fixations of forming the first ‘gist’ of the image before directing the gaze to the semantically salient object.

#### Language modelling

To capture how narrative context influences visual sampling, we used language modelling as a way to quantify the semantic salience of objects from the high-level contextual representation of the story. It should be stressed that these language-derived narratives were obtained completely separately (by different narrators) and that our observers were not exposed to language in any way—they were simply viewing the images. Yet, the differences in sampling behaviour between meaningful and shuffled contexts were well explained by this language-derived model. This suggests that language is an effective expression of the structure of narrative meaning, paralleling the representations participants naturally aimed to construct while simply viewing the images. More broadly, this approach highlights a promising link between language-based representation of meaning and visual sampling, suggesting that language can serve as a powerful tool for modelling top-down influences of cognition on visual processing, and emphasizes the cross-links between language and vision.^45^

The cross-link between vision and language has been, especially in the recent years, recognized increasingly, with more studies linking visual activity or eye movements to cognition via language modelling.^46,47^ For instance, it has been shown that by embeddings of a large language model to scene captions can account for responses in ventral visual regions during natural scene viewing.^46^ Similarly, it has been shown that using language models to describe how all objects in scene images relate to on another can explain what objects are fixated on the most.^24^ Note that these studies have still focused on single images and thus used language to quantify visual elements in a rather non-contextualized way, whereas we have here used a contextualized model that draws predictions over a longer timeframe. A useful direction forward will be to combine these somewhat still separate fields – on visual and scene perception, computational psycholinguistics, narrative viewing to study how language may offer a useful representational format that accounts for cognitive influences on visual perception and sampling.

#### Visual salience

Our results also underscore the importance of investigating visual sampling in ecologically valid ways that reflect how visual input is encountered in the real world. Namely, while visual salience models like DeepGaze-II^4^ and GBVS^6^ powerfully predict gaze patterns from static features, they are inherently lacking any concept of temporal context. Real-world visual experience, in contrast, is always embedded in rich temporal context and follows a coherent structure: we rarely encounter completely unrelated and random snapshots one after the other. Instead, visual input is embedded in a broader context that changes in a temporally predictable way, in accordance with our internal models. These findings point to the need for future models to incorporate rich contextual and semantic structure, and to move beyond single-image paradigms towards more ecologically valid settings.^48–50^

It is important to mention that visual salience *did* explain a range of sampling behaviour— the most visually salient objects (as defined by DeepGaze-II) were fixated both earlier and more frequently than the semantically salient ones. This confirms that bottom-up image-computable features exert a strong influence on gaze behaviour. However, it should also be noted that this was much more pronounced for DeepGaze-II, and not for GBVS, another visual salience model, which relies purely on image-computable features. This difference likely stems from DeepGaze-II being trained on human fixations and/or the use of deep neural network as its basic architecture for extracting features of salience. Most importantly, the context-dependent modulation of gaze behaviour was observed across both models, further reinforcing the robustness of the findings.

We observed that in terms of temporal evolution, the first few fixations were much more likely to be directed at the most visually salient object. This is in line with prior investigations of attentional deployment over time^44^, which have also found that early fixations are directed to the more physically (or visually) salient aspects of the image, regardless of the task, whereas later fixations are less driven by the visual salience.^40^ Still, we observed also an early diminished influence of visual salience when images were presented in the temporally coherent order, suggesting that task (or goals) can act already early on.

It is important also to relate the findings of visual salience to a simpler mechanism, central fixation bias, which is inherently present in both models we investigated – DeepGaze-II and GBVS. We observed that individuals were also more likely to fixate on more central objects for the first two fixations than afterwards, but this was no different depending on the temporal order in which images were presented (coherent or shuffled). However, when accounting for sampling frequency and latency of all objects, then even the central fixation model accounted for some differences between coherent and shuffled orders. Nonetheless, both GBVS and DeepGaze-II’s Spearman’s correlation values were much higher than for central fixation bias model, suggesting that other aspects of the models than purely object’s distance to the centre still accounted for sampling latency and frequency.

### Limitations

There are several limitations that we need to discuss with our work. Our semantic salience metric used here can be seen as a proof of principle of how such a metric could be obtained and related to eye movements. However, it is not image-computable yet, and currently relies on several manual processing stages (such as manual assignment of phrases to objects). We believe that with ever-increasing advancements in the field, this is very likely become more automatized and scale up, including to dynamic settings of movie watching (with for instance using automatic Gemini caption system to obtain semantic embeddings of ongoing movie development). This will also allow to test several previously stated predictions, for instance that with dynamic stimuli more of gaze behaviour is driven by visual, as opposed to semantic (or cognitive) influences (^17^, some evidence in this direction^47^).

Secondly, the stimuli we used were cartoon-like images that were hand drawn, which presents a limitation for the visual models we used. Namely, the convolutional neural network VGG-19^35^, which forms the basis for DeepGaze-II, was trained on photographic images, and thus does not perform very well in classifying cartoon-like images.^51^ Therefore we acknowledge that the performance of DeepGaze-II here was likely worse than it would have been on photographic stimuli, or if it would have been additionally fine-tuned on hand-drawn illustrations. Furthermore, we also averaged all the salience values within the object, thus possibly further limiting the power of DeepGaze-II which intrinsically predicts fixations to a higher resolution.

#### Conclusion and future outlook

The big open question for the future work is how exactly cognition influences eye movements, and how different types of salience interact with each other and task demands to shape gaze behaviour. Here we have primarily elucidated *that* semantic salience influences gaze behaviour and how it could be computationally quantified, but not addressed the mechanistic question. Based on the obtained results, we would speculate that cognitive influences on eye sampling behaviour fall within the ‘second wave’ of influence after the first feedforward sweep of more physical, or visual salience. Thus, after the gist of the image is depicted, then in combination with ongoing forming model of the world, more specific ‘guesses’ and directions towards semantic salience to further improve the narrative model can be drawn. We have focused here primarily on surprisal as the important metric for model update, but it could be that also other factors have additional, or different influences – such as stimulus importance, relevance, entropy etc. Moreover, while we have here formed a group model of semantic salience, one further open question for the future is to explore how a model of each viewer develops on the go; and how every single fixation adds to the forming model of the narrative. One possible way to better understand how individuals build a model across images, may be by employing the ‘thinking aloud’ paradigm^52^ where after every image has been presented, individuals would report what they have seen. In this manner, one would also have a better insight into one individual’s forming model of the events. Being able to quantify the informational update per fixation and within the observer, could open up a whole array of further personalized questions to address, including moving from understanding typical visual sampling into clinical domains. In doing so, also more eye movement metrics should be explored, beyond the ones included here – such as temporal evolution (i.e. scanpath), and exploring processing also outside of fovea into periphery^53^ to embrace the richness of gaze behaviour.

To conclude, we have here introduced a novel approach for modelling meaning in the visual world through language, and successfully linked it to gaze behaviour. Our study can be seen as a testbed for further exciting applications that can build on this framework. With continuously evolving advances in AI, this approach could be scaled up to dynamic and immersive settings, such as film viewing or real-world navigation. Modelling how the cognitive and visual salience jointly guide attention moment-by-moment may yield deeper insights into how we sample our environments, make sense of it, and when we disengage. Ultimately, our findings reveal that eye movements are not just a window into what we see, but also into how we understand the bigger picture.

## Data and code availability

All materials, data, and code related to the current manuscript are openly available (upon user registration) on a Radboud Data Repository Site (Eye movements during narrative viewing; https://doi.org/10.34973/014c-t415).

## Acknowledgements

EB was supported by European Union’s Horizon Europe research and innovation programme under the Marie Skłodowska-Curie grant agreement for individual fellowship (101106569). LBS was supported by a by European Union’s Horizon Europe research and innovation programme under the Marie Skłodowska-Curie grant agreement for individual fellowship (101111402). CHH was supported by the Italian Ministry of University and Research (MUR) within the PNNR NextGenerationEU program (No. 0000027).

## Author contributions

EB: conceptualization, methodology, investigation, formal analysis, visualization, writing – original draft, review and editing.

LMS: formal analysis (supporting), writing – review and editing. CHH: methodology (supporting), formal analysis (supporting), writing – review and editing. MVP: conceptualization (supporting), writing – review and editing. FPdL: conceptualization, formal analysis (supporting), visualization (supporting), writing – review and editing.

## Supplementary figures

**Supplementary Figure 1.**
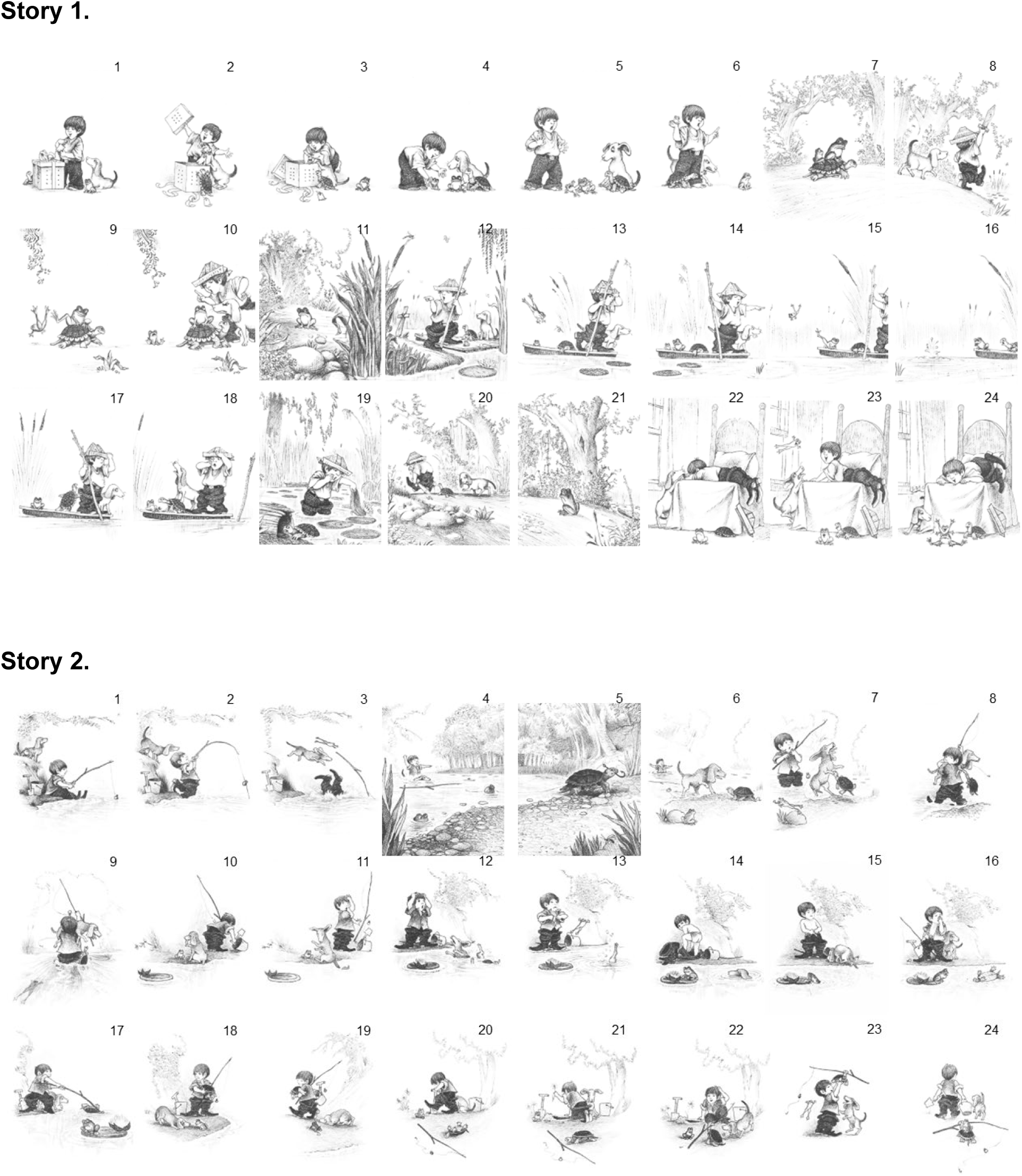
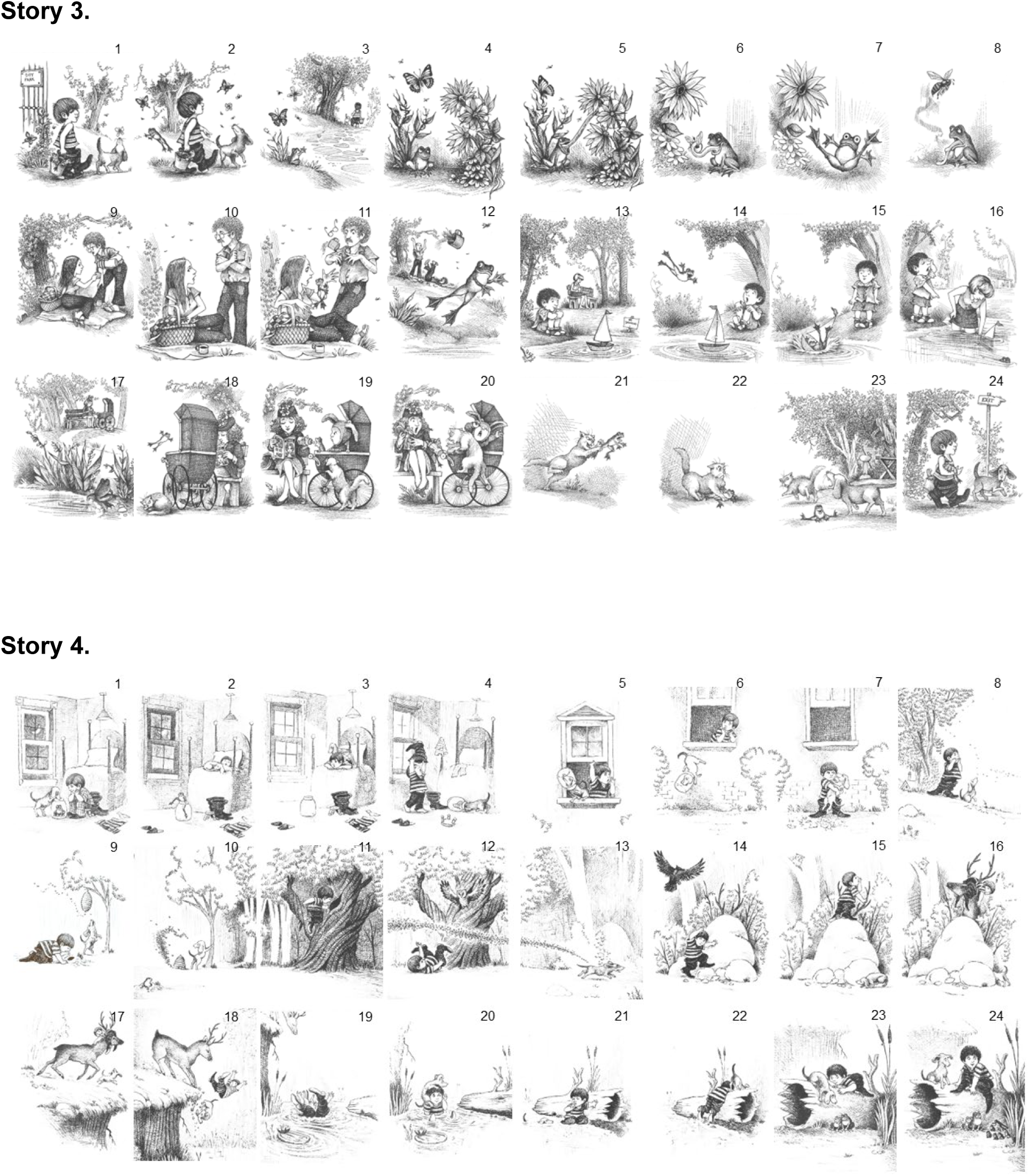
Images from all of the four stories, as presented in the coherent condition. Note that the order of presentation (story 1-4) was randomized between participants.

**Supplementary Figure 2.**
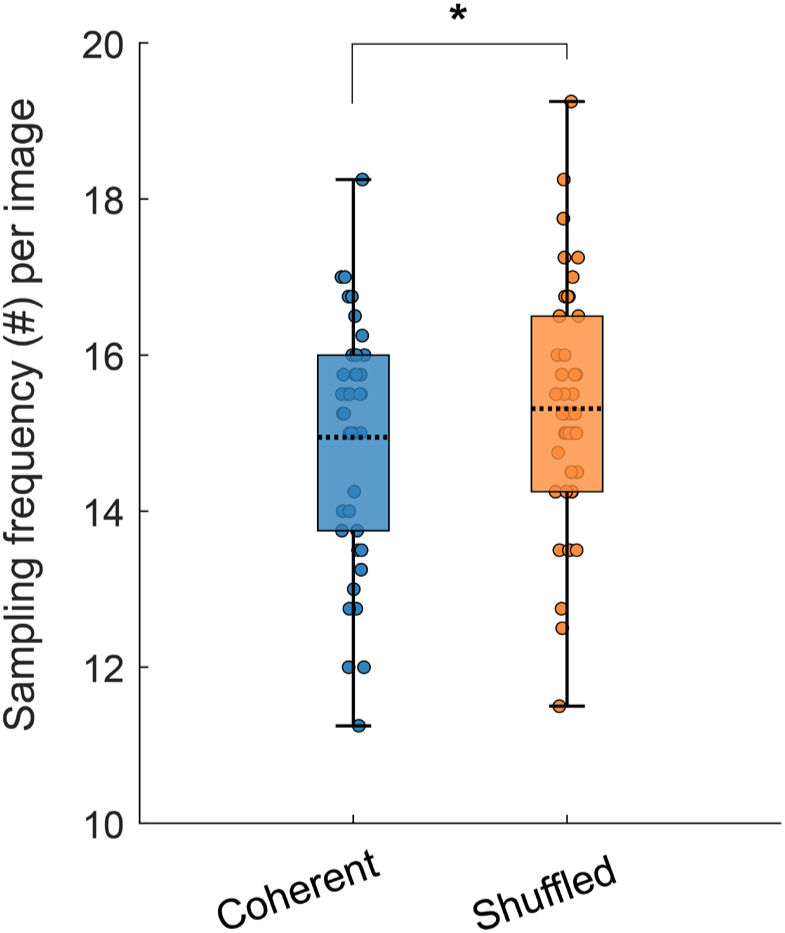
Fewer fixations during presentation in the coherent than shuffled order. Participants made on average fewer fixations during the 5-second image presentation when images were presented in coherent than shuffled condition (*t*_(41)_=2.84, *p*=6.94e^-3^).

**Supplementary Figure 3.**
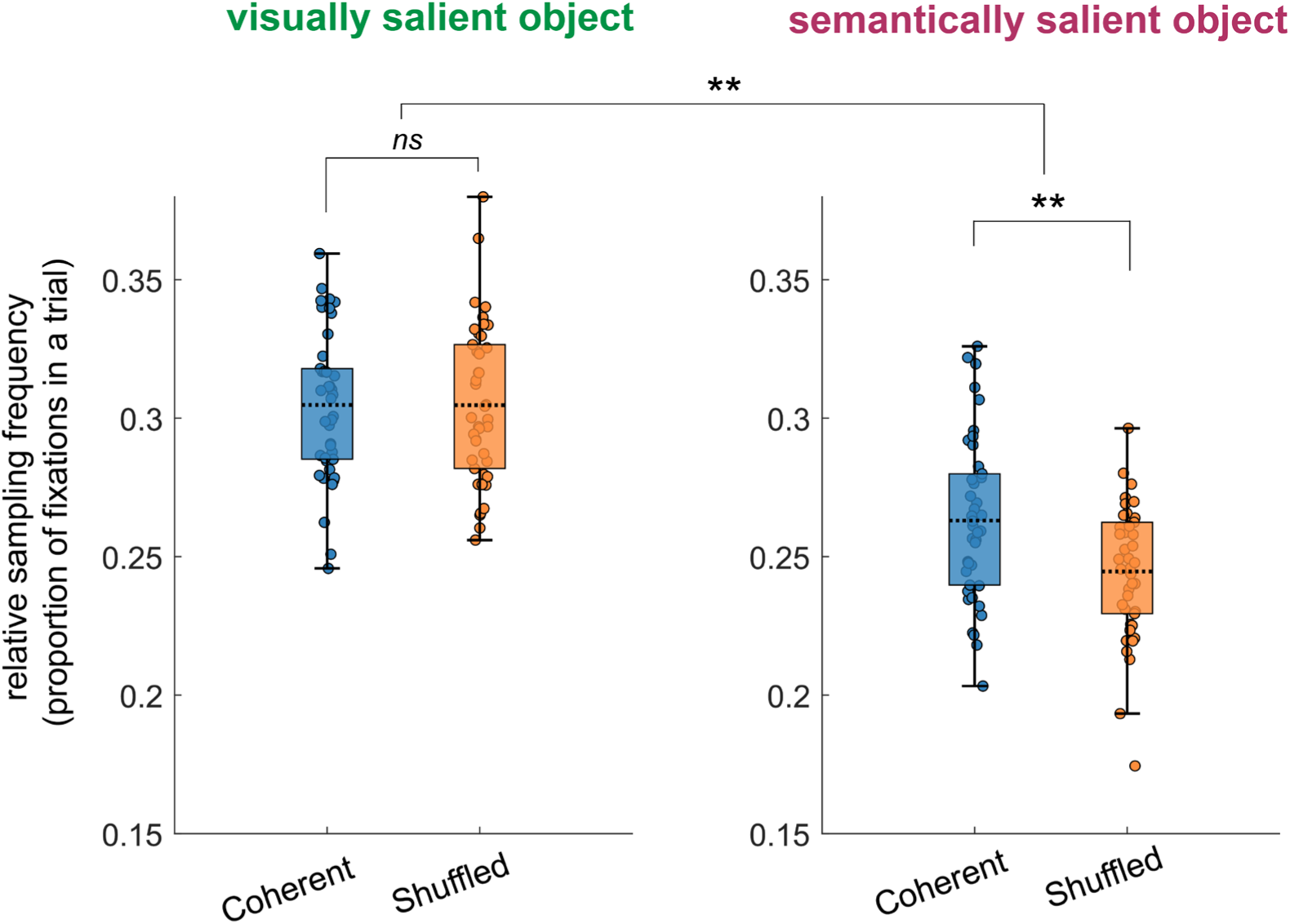
Relative frequency of fixations to the most visually and semantically salient object, split by condition (coherent or shuffled order). The relative frequency was calculated by dividing the number of fixations to the most visually or semantically salient object by the absolute number of fixations in each individual trial. Also when considering the proportion of fixations that were directed to the most salient object, there is a condition-dependent modulation of saliency on sampling behaviour (interaction of order x salience type *F*_(1,41)_ = 10.212, *p*=.0027). Post-hoc t-tests revealed that a higher proportion of fixations were directed to the most semantically salient object when in coherent than shuffled order (*t*_(41)_ = 3.2, *p*=2.60e^-3^). There was no difference in the relative proportion of fixations directed at the most visually salient object depending on the order (*t*_(41)_<1). In the figure ** indicates significance level of *p*<.01 and *ns* means not significant (*p*>.05).

**Supplementary Figure 4.**
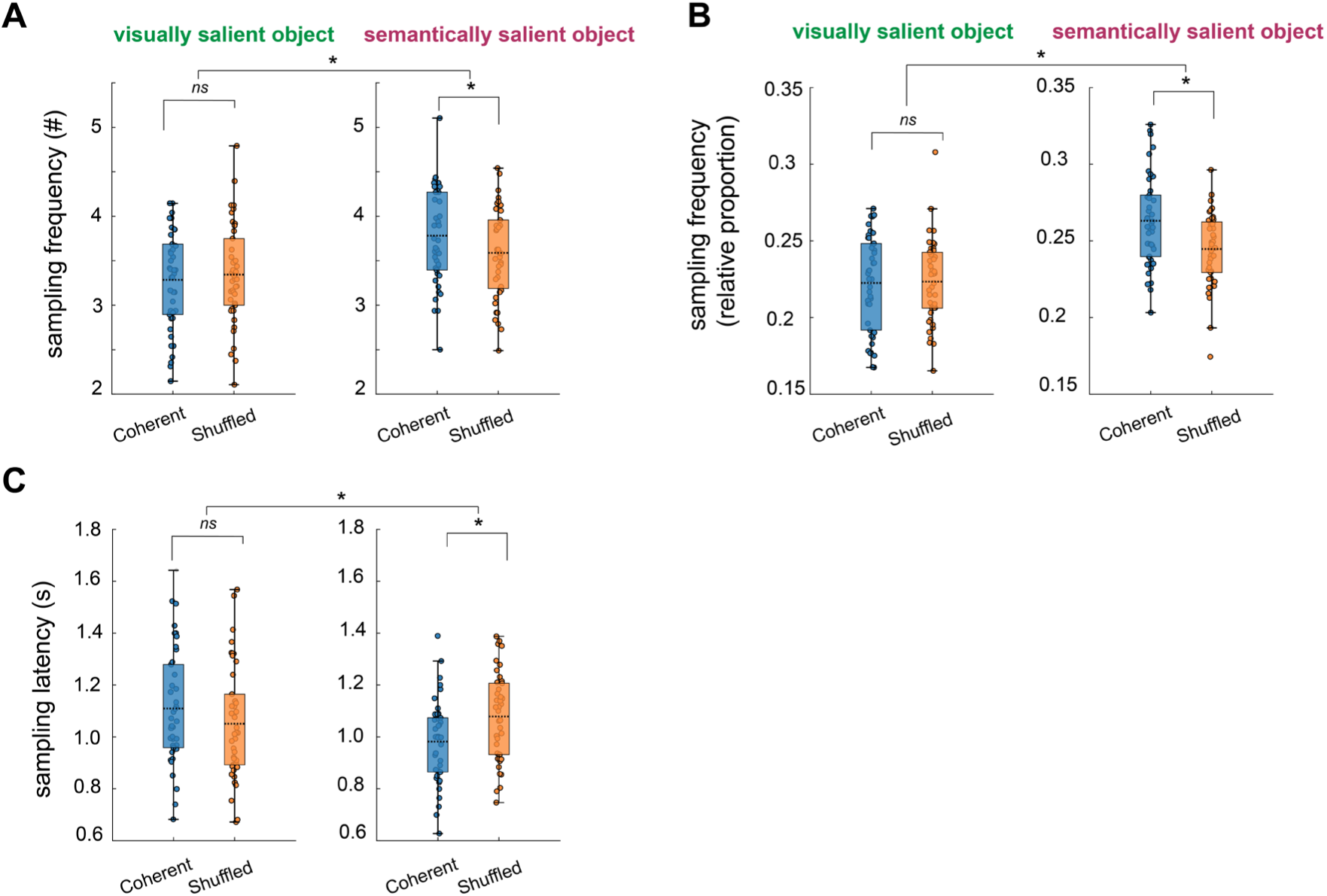
Sampling frequency and latency for the most salient objects using a different salience model, Graph-Based Visual Salience (GBVS). Results are comparable to the most salient object as defined by DeepGaze-II. **A)** Fixation frequency (absolute fixation number) on the most salient objects, with the most visually salient being defined by salience model GBVS. The frequency of fixation depended on the interaction between the salience type and order of presentation (coherent vs. shuffled) (ANOVA: *F*_(1,41)_=6.04, *p*=.018). There was also a main effect of model type, with semantically most salient object fixated more often than the GBVS-defined most visually salient object (*F*_(1,41)_=39.87; *p*=1.54e^-7^). No main effect of order was observed (*F*_(1,41)_=0.70, *p*=.41). Post-hoc t-tests on the effect of context condition for the visually most salient object revealed no difference (*t*_(41)_=0.52, *p*=.61). In the figure * indicates significance level of *p*<.05 and *ns* means not significant (*p*>.05). **B)** When fixation frequency was quantified as a relative, as opposed to absolute, fixation count, the results were similar. ANOVA interaction of order x salience type: *F*_(1,41)_=6.44, *p*=.015, main effect of salience type: *F*_(1,41)_=55.91, *p*=3.52e^-9^, main effect of order: *F*_(1,41)_=2.552, *p*=.117. For post-hoc t-test there was no significant difference between conditions for sampling of the visually most salient object (*t*_(41)_=0.11, *p*=.91). **C)** For sampling latency, similar patterns of results were observed when defining the most salient object with GBVS instead of DeepGaze-II. We observed an order-specific modulation of the salience type (interaction: *F*_(1,41)_=6.652, *p*=.0136), but no significant main effect of either salience type (*F*_(1,41)_=2.90, *p*=.096), or order (*F*_(1,41)_=0.36, *p*=.55). The post-hoc t-test revealed no significant difference between conditions in terms of latency to the most visually salient object (*t*_(41)_=1.144, *p*=.26).

**Supplementary Figure 5.**
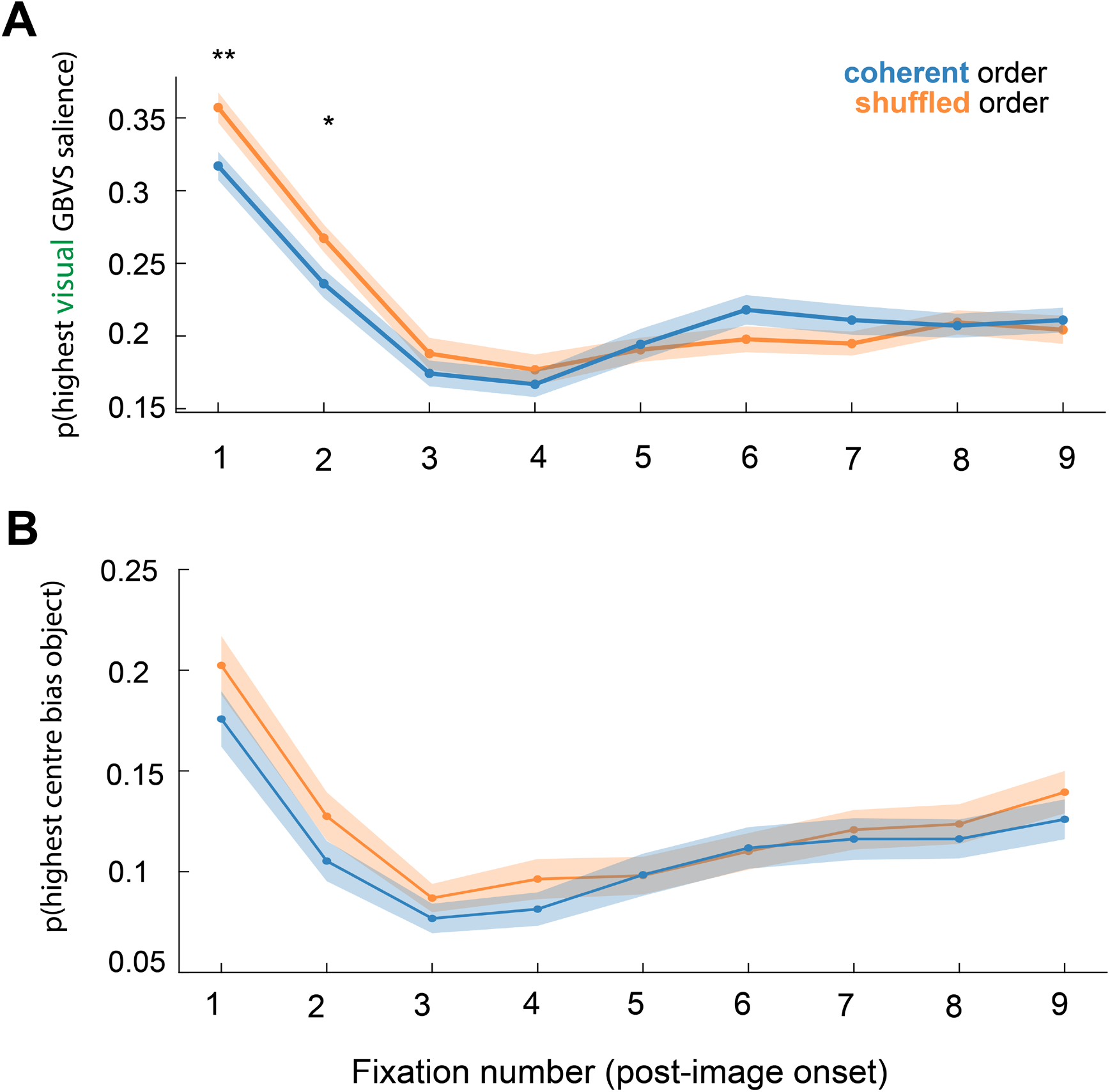
Likelihood of sampling the most visually salient object, as defined by GBVS, or object highest in terms of central bias tendency. Fixation numbers denote numbers from the start of image onset (1 being the first fixation). **A)** The likelihood of sampling the most visually salient object as defined by GBVS over time is similar as was observed for DeepGaze-II, with first few fixations being more likely to be directed at the most visually salient object when images are presented in the shuffled than coherent temporal order (fixation 1: *t*_(41)_=2.79, *p*=.008 and fixation 2: *t*_(41)_=2.46, *p*=.018). **B)** The likelihood of sampling the object scoring the highest in terms of central tendency also diminished from first few fixations onwards. There was no difference in likelihood of fixation towards the centre depending on order (coherent vs. shuffled order: ANOVA interaction fixation number x context: *F*_(8,328)_=0.53, *p*=.88).

**Supplementary Figure 6.**
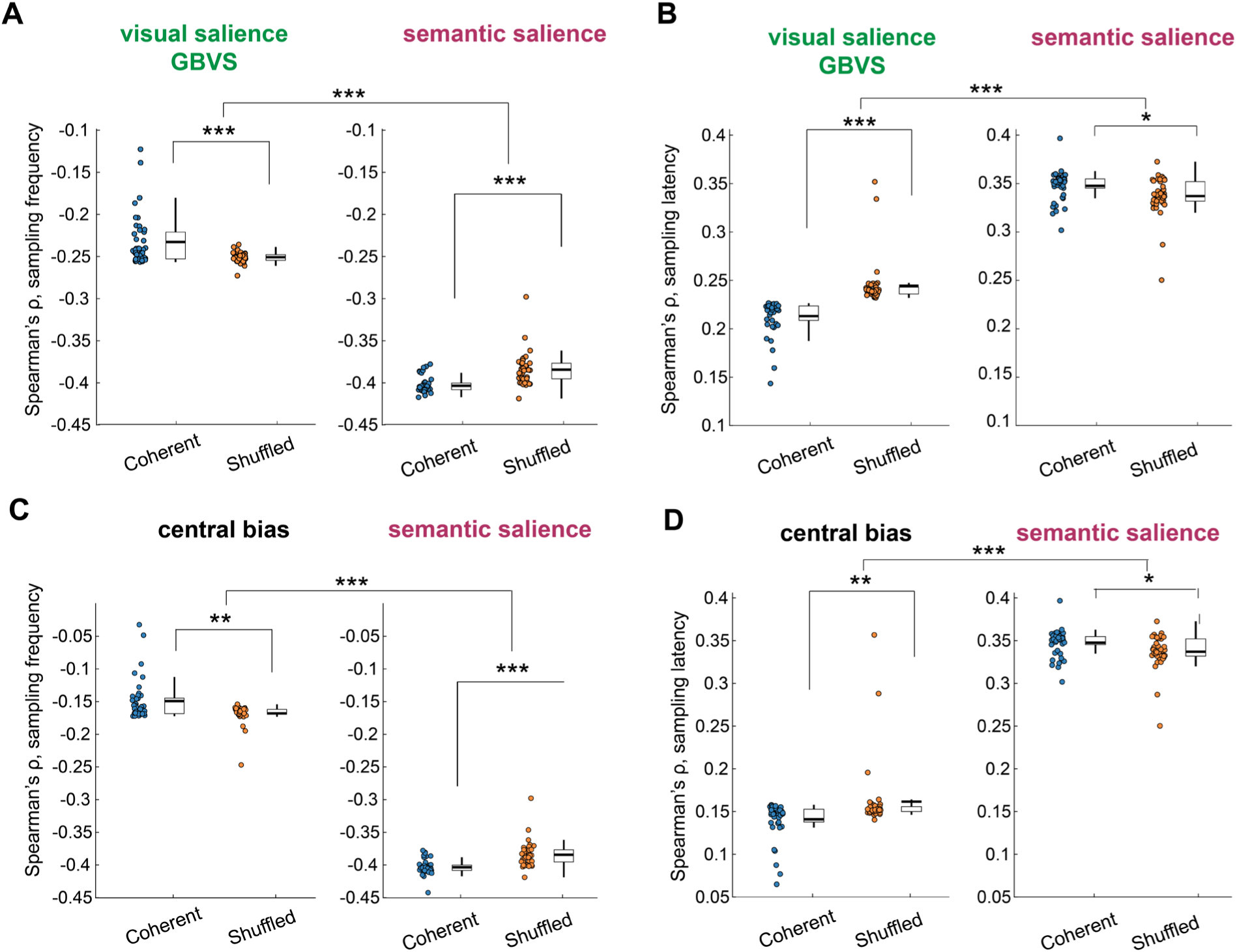
Spearman correlation, examining sampling behaviour across all objects with a different model of visual salience, GBVS, and the central bias tendency model. **A)** Sampling frequency. The interaction of salience model (GBVS vs. semantic salience) with condition order was significant (*F*_(1,41)_=30.09, *p*=2.327e^-6^), and also post-hoc tests for visual salience, as defined with GBVS (*t*_(41)_=3.65, *p*=7.45e^-4^). **B)** Sampling frequency. The interaction of model (GBVS visual salience vs. semantic salience) with condition order was significant (*F*_(1,41)_=27.61, *p*=4.92e^-6^), and also post-hoc tests for visual salience, as defined with GBVS (*t*_(41)_=5.63, *p*=1.5e^-6^). In the figure * indicates significance level of *p*<.05, ** significance level of *p*<0.01 and *\*\*\** significance level of *p*<0.001.**C)** Sampling frequency. The interaction of model type (centre bias or semantic salience model) with condition order was significant (*F*_(1,41)_=19.56, *p*=7.01e^-5^), and also post-hoc tests for the central bias model, with the sampling frequency in the shuffled order having a stronger (negative) Spearman correlation than sampling when the order was coherent (*t*_(41)_=3.05, *p*=4.0e^-3^). **D)** Sampling latency. The interaction of model type with condition order was also significant (*F*_(1,41)_=7.721, *p*=8.23e^-3^), and also post-hoc tests for the central bias model, with the sampling frequency in the shuffled order having a higher Spearman correlation than sampling when the order was coherent (*t*_(41)_=2.42, *p*=2.0e^-3^).

